# Pan-cancer analysis of RNA binding proteins (RBPs) reveals the involvement of the core pre-mRNA splicing and translation machinery in tumorigenesis

**DOI:** 10.1101/341263

**Authors:** Bin Zhang, Kamesh R. Babu, Chun You Lim, Zhi Hao Kwok, Jia Li, Siqin Zhou, Henry Yang, Yvonne Tay

**Affiliations:** Cancer Science Institute of Singapore, National University of Singapore, Singapore 117599, Singapore; Department of Biochemistry, Yong Loo Lin School of Medicine, National University of Singapore, Singapore 117597, Singapore

## Abstract

RNA binding proteins (RBPs) are key regulators of posttranscriptional processes such as RNA maturation, transport, localization, turnover and translation. Despite their dysregulation in various diseases including cancer, the landscape of RBP expression and regulatory mechanisms in human cancer has not been well characterized. Here, we analyzed mRNA expression of 1487 RBPs in ~6700 clinical samples across 16 human cancer types and found that there were significantly more upregulated RBPs than downregulated ones in tumors when compared to their adjacent normal tissues. Across almost all of the 16 cancer types, 109 RBPs were consistently upregulated (cuRBPs) while only 41 RBPs were consistently downregulated (cdRBPs). Integrating expression with the copy number and DNA methylation data, we found that the overexpression of cuRBPs is largely associated with the amplification of copy number, whereas the downregulation of cdRBPs may be a result of epigenetic silencing mediated by DNA methylation. Furthermore, our results indicated that cuRBPs could work together to promote cancer progression potentially through the involvement of splicing and translation machinery, while cdRBPs might function independently to suppress tumorigenesis. Additionally, we focused on colon cancer and identified several novel potential oncogenic RBPs, such as PABPC1L which might promote cancer development via regulating the core splicing machinery. In summary, we showed distinct expression landscapes, regulatory mechanisms and characteristics of cuRBPs and cdRBPs and implicated several novel RBPs in cancer pathogenesis. Moreover, our results suggest that the involvement of the core pre-mRNA splicing and translation machinery could be critical in tumorigenesis.

## Introduction

RNA-binding proteins (RBPs) are a group of conserved proteins in eukaryotes, which play essential roles in co-transcriptional and posttranscriptional gene regulation, including RNA maturation, RNA turnover, RNA localization and translation (Glisovic et al. 2008). RBPs can interact with RNAs to form protein-RNA complexes, such as ribosomes that serve as the basic translational machinery, and small nuclear ribonucleo proteins (snRNPs) which are core premRNA splicing machinery. RBPs can also regulate splicing, translation by controlling accessibility and activity of the basic machineries, for instance the splicing enhancer/repressor and translation regulators (Moore and Proudfoot 2009; Fu and Ares 2014). Based on RNA binding domain prediction and manually selection of RBPs from literature, a census of 1542 human RBPs has been established in recent years (Gerstberger et al. 2014). However, only a small proportion of these RBPs have been functionally characterized.

Since RBPs have diverse functions in posttranscriptional gene regulation, their appropriate expression is critical to many biological processes, such as cell differentiation, proliferation and cell fate transition. For example, the RBP genes MBNL1 and MBNL2 control splicing of the 18th exon of transcription factor FOXP1 to switch pluripotency and reprogramming of embryonic stem cells (ESCs), while the RBP gene NUDT21 directs alternative polyadenylation of thousands of transcripts to control cell fate transition of ESCs (Gabut et al. 2011; Han et al. 2013; Brumbaugh et al. 2018). Dysregulation of RBPs could cause severe human diseases, including cancer (Yeo 2014). Based on somatic mutation analysis from more than 3000 cancer patients, 31 splicing factors have been predicted to be cancer drivers (Sveen et al. 2016). Among them, SF3B1 is mutated in around 10-15% of chronic lymphocytic leukemia (CLL) patients and is associated with poor survival rate (Quesada et al. 2011; Wang et al. 2011). In addition, mutations within four RBP genes U2AF, ZRSR2, SRSF2 and SF3B1 were very frequent (~70%) in a cohort of myelodysplasia (MDS) patients (Yoshida et al. 2011). Besides mutation, it has been reported that dysregulation of core splicing and translational machinery gene expression is essential for MYC-mediated lymphomagenesis (Barna et al. 2008; Koh et al. 2015). Moreover, it has been revealed that the RBP gene NELFE could promote cancer progression via selectively regulating MYC-associated genes (Dang et al. 2017).

Despite extensive studies on selected RBPs, the general role of RBPs in cancer development is still not clear. In normal human tissues, it has been reported that the expression of RBPs is significantly higher than that of transcription factors (TFs), other protein coding genes (PCGs), miRNAs and lincRNAs (Gerstberger et al. 2014; Kechavarzi and Janga 2014). Furthermore, ribosomal proteins showed distinct expression signatures between tumor and normal tissues (Guimaraes and Zavolan 2016). However, the global expression changes of RBPs in cancers remain elusive. One recent study found that RBPs tend to be predominantly downregulated in tumors (Wang et al. 2018). In contrast, another study revealed that RBPs tend to be upregulated in hepatocellular carcinoma (HCC) comparing with normal liver tissues (Dang et al. 2017). Therefore, there is a pressing need to study the overall expression landscape of RBPs in cancers, understand the regulatory mechanisms underlying dysregulation and investigate their roles in cancer development.

To address these questions, we analyzed the mRNA expression of 1487 RBPs in ~ 6700 clinical samples across 16 human cancer types in The Cancer Genome Atlas (TCGA) project. Our results revealed that there are significantly more RBPs showing upregulation in tumors compared to their adjacent normal tissues. In almost all of the 16 cancer types we studied, 109 RBPs are consistently upregulated while only 41 RBPs are consistently downregulated. Next we integrated expression with somatic copy number alteration (SCNA) and DNA methylation data from the same clinical samples, and found that upregulation of consistently upregulated RBPs (cuRBPs) in tumors are largely associated with amplification of copy number, whereas downregulation of the consistently downregulated RBPs (cdRBPs) is most likely a result of epigenetic silencing mediated by DNA methylation. Utilizing the gene sets from the molecular signature database, we also found several TFs and miRNAs, such as MYC, MAZ, SP1, E2F1, miR-25 and miR-19a, that could be responsible for the dysregulation of several cuRBPs and cdRBPs in tumors. Furthermore, based on our integrated analysis of **tu**mor **s**uppressor and **on**cogene (TUSON) explorer, protein-protein interaction (PPI) network, protein complex and survival analysis we suggest that cuRBPs might function together to promote cancer progression through the involvement of the core pre-mRNA splicing and translation machinery. In contrast, cdRBPs might function independently to suppress tumorigenesis. Specifically, in colon adenocarcinoma (COAD), we identified several potential oncogenic RBP candidates, such as PABPC1L which might promote cancer progression in COAD by regulating the core pre-mRNA splicing machinery. Taken together, our study provides insights to understand the dysregulation of RBPs during tumorigenesis and identifies novel cancer-related RBPs which might be helpful for clinical applications. Moreover, our results also suggested that the dysregulation of core spliceosome and ribosome biogenesis could be critical for cancer progression.

## Results

### RBPs tend to be upregulated in tumors compared with normal tissues

To dissect the dysregulation of RBPs in human cancers, we analyzed mRNA expression of 1478 RBPs in ~6700 clinical samples across 16 cancer types based on RNA sequencing data from TCGA (Supplemental Table S1). The other 17 cancer types were excluded due to lack of sufficient normal samples (n < 10). For each cancer type, dysregulated RBP genes were identified by comparing the expression in tumors to their adjacent normal tissue samples based on the same criteria (BH-adjusted *p* < 0.001) as the previous study by Dang *et al*. Interestingly, there were many more upregulated RBPs than downregulated RBPs in 14 out of the 16 cancer types we studied, while this trend is not observed for TF genes (Figure 1A). This is consistent with a previous study, which reported predominant upregulation of RBPs in HCC (Dang et al. 2017). Besides the numbers of up and down regulated RBPs, we also analyzed the global expression changes of all RBPs and observed a general upregulation trend in almost all of cancer types, except kidney chromophobe (KICH) and thyroid carcinoma (THCA) (Supplemental Fig. S1).

**Figure 1:**
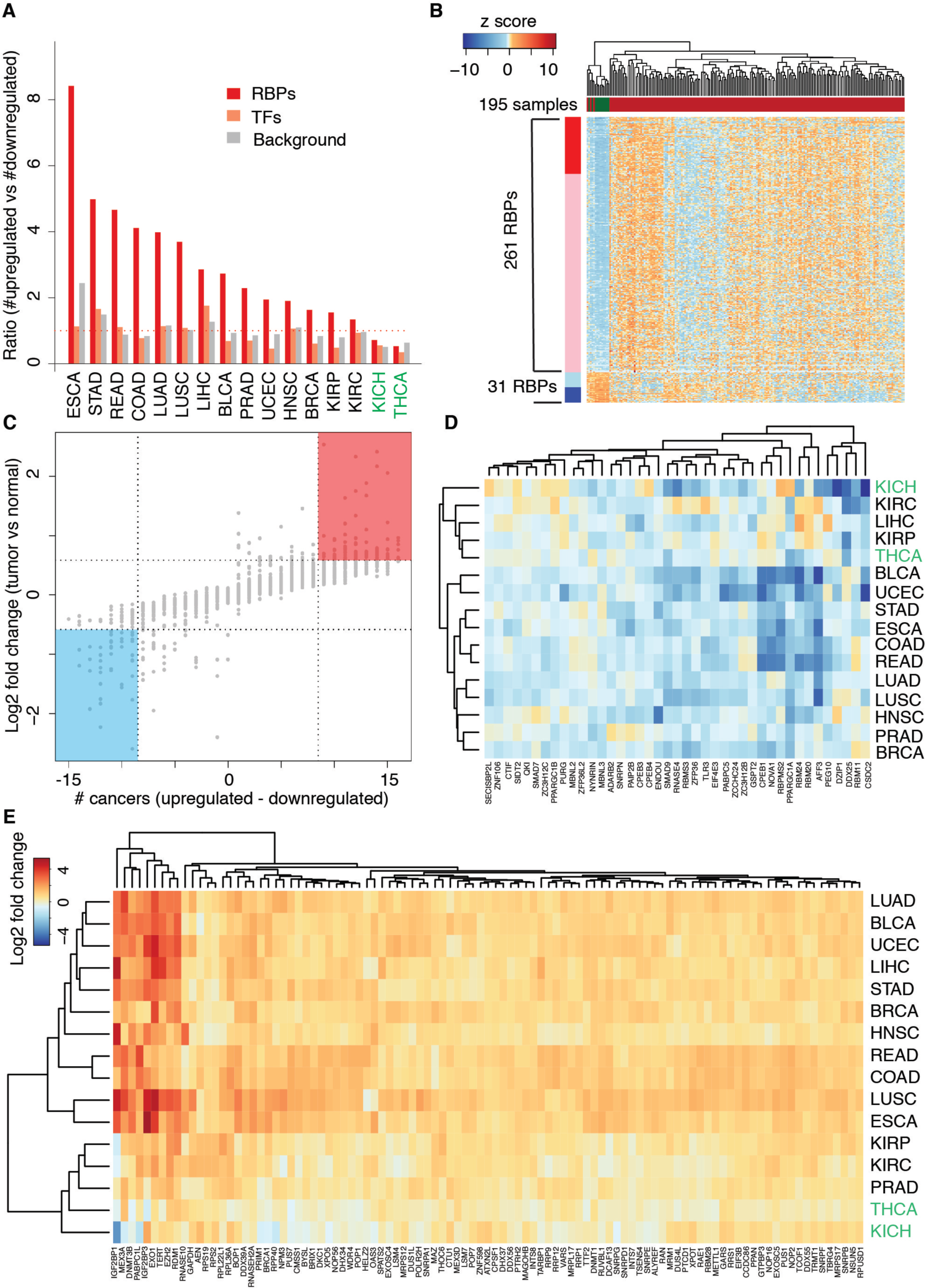
Expression profiles of RBPs in different human cancer types. (A) The ratio of the number of upregulated genes to that of downregulated ones when comparing tumor with normal samples across different cancer types. Background: all PCGs. (B) Heatmap illustrating the expression of significantly differentially expressed RBPs between tumor and normal in ESCA (BH-adjusted *p* < 0.001). The left bar indicates the upregulated and downregulated RBP genes (red: log_2_ fold change > 1; pink: log_2_ fold change > 0 and < 1; light blue: log_2_ fold change < 0 and > -1; dark blue: log_2_ fold change < -1). The indicates upper bar the tumor and normal samples (red: tumor, green: normal). (C) The directionality and amplitude of the expression change of RBPs in tumor compared with normal. The X axis represent the directionality that is defined by the number of cancer type RBPs show upregulated expression minus the number of cancer types where RBPs show downregulated expression. The Y axis is the amplitude of the expression change in terms of the average of the fold change across 16 cancer types. (D) and (E) Heatmaps illustrating the fold change of 41 cdRBPs (D) and 109 cuRBPs (E) across 16 cancer types.

Next, we analyzed the expression profile of 292 significantly dysregulated RBPs in 195 samples of esophageal carcinoma (TCGA-ESCA) in greater detail since it showed the most striking upregulation bias. The hierarchical clustering of different samples based on the expression was able to separate the tumor from normal. Moreover, it was apparent that RBPs were predominantly upregulated rather than downregulated in tumors (261 upregulated versus 31 downregulated) and such bias was not affected by more stringent criteria with fold change of 2 and adjusted *p* of 0.001 (60 upregulated versus 15 downregulated) (Figure 1B). We also examined the distribution of fold changes of all dysregulated RBPs and found that this trend was not affected by different cutoffs (Supplemental Fig. S2). In bladder urothelial cancer (BLCA), however if such criteria are required, despite we could observed a general upregulation of RBPs, the number of upregulated ones is slightly lower than that of downregulated (Supplemental Fig. S3). This was the only instance in which we found predominant down regulation of RBPs expression in cancers as previous report (Wang et al. 2018), whereas our results suggested that in general RBPs tend to be upregulated in most human cancer types we studied. To further confirm this biased expression pattern, we analyzed mRNA expression in the independent GEO public dataset GSE102083 which contains microarray data from 152 HCC and 91 normal liver samples. Consistently, we still observed that the number of upregulated RBPs is many more than that of downregulated ones in tumor samples when compared to normal (Supplemental Table S2).

As RBPs could be dysregulated across different cancer types, we examined two features for each RBP: 1) difference in the number of cancer types of upregulation versus downregulation; 2) log2 fold change tumor/normal. Based on this we found 109 cuRBPs and 41 cdRBPs (Material and Methods) (Figure 1C). This number suggested that RBPs tend to be upregulated in tumor compared to normal tissues. For the 41 cdRBPs, CPEB1, NOVA1, RBPMS2, PPARGC1A, RBM20, RBM24 and AFF3 exhibited drastic downregulation pattern in many cancer types (Figure 1D). The CPEB family proteins has been implicated in translation control of cancer cells, while NOVA1 has been found to be downregulated in the gastric cancer microenvironment (Fernandez-Miranda and Mendez 2012; D’Ambrogio et al. 2013; Yoon et al. 2016). Among 109 cuRBPs, IGF2BP1, MEX3A, DNMT3B, PABPC1L, IGF2BP3, EXO1, TERT, EZH2 and RDM1 showed the most striking overexpression pattern (Figure 1E). DNMT3B and EZH2 are well known epigenetic regulators and both have strong oncogenic functions, while the promoter of TERT is recurrently mutated in many cancers (Beaulieu et al. 2002; Rhee et al. 2002; Varambally et al. 2002; Simon and Lange 2008; Huang et al. 2013; Vinagre et al. 2013). IGF2BP1 and IGF2BP3 are two oncofetal proteins which have been reported to promote adhesion, migration and invasiveness of tumor cells by mediating mRNA stability and translation (Bell et al. 2013; Lederer et al. 2014). However, the functional roles of PABPC1L, MEX3A and EXO1 in tumorigenesis remain totally unexplored. Taken together, our results revealed that RBPs tend to be upregulated in human cancers and identified many novel RBPs that may play important but hitherto uncharacterized roles in cancer.

### Dysregulation of cuRBPs is associated with amplification of copy number while dysregulation of cdRBPs could be due to epigenetic silencing

Gene expression can be modulated by multiple mechanisms, such as copy number alteration, the abundance of transcription factors and miRNAs, and DNA methylation at the promoter region (Stranger et al. 2007; Hobert 2008; Jones 2012). To explore their relationship with dysregulation of RBP expression in cancer, we first analyzed somatic copy number alteration (SCNA) by comparing the tumor to normal samples. As expected, the cuRBPs have significant amplification of copy numbers (Mann-Whitney test, p < 4.9e-77, 16 cancer types together), while the cdRBPs have significant loss of copy numbers (Mann-Whitney test, p < 6.7e-22, 16 cancer types together). This trend is consistent in almost all analyzed cancer types except KICH and THCA (Figure 2A). To investigate whether the SCNA do result in the dysregulation of RBPs in cancer, we analyzed the correlation between SCNA and mRNA expression for each gene. Indeed, there is a general positive correlation between expression and SCNA for all the PCGs, cdRBPs, consistently upregulated TFs (cuTFs) and consistently downregulated TFs (cdTFs). Interestingly, we observed a much stronger positive correlation for the cuRBPs (median of *r* = 0.42, *p* < 2.1e-160, Mann-Whitney test) compared to the PCGs background (median of *r* = 0.2) (Figure 2B). This suggests that the overexpression of cuRBPs in tumors is caused by amplification of copy numbers, which might be critical to drive cancer progression.

**Figure 2:**
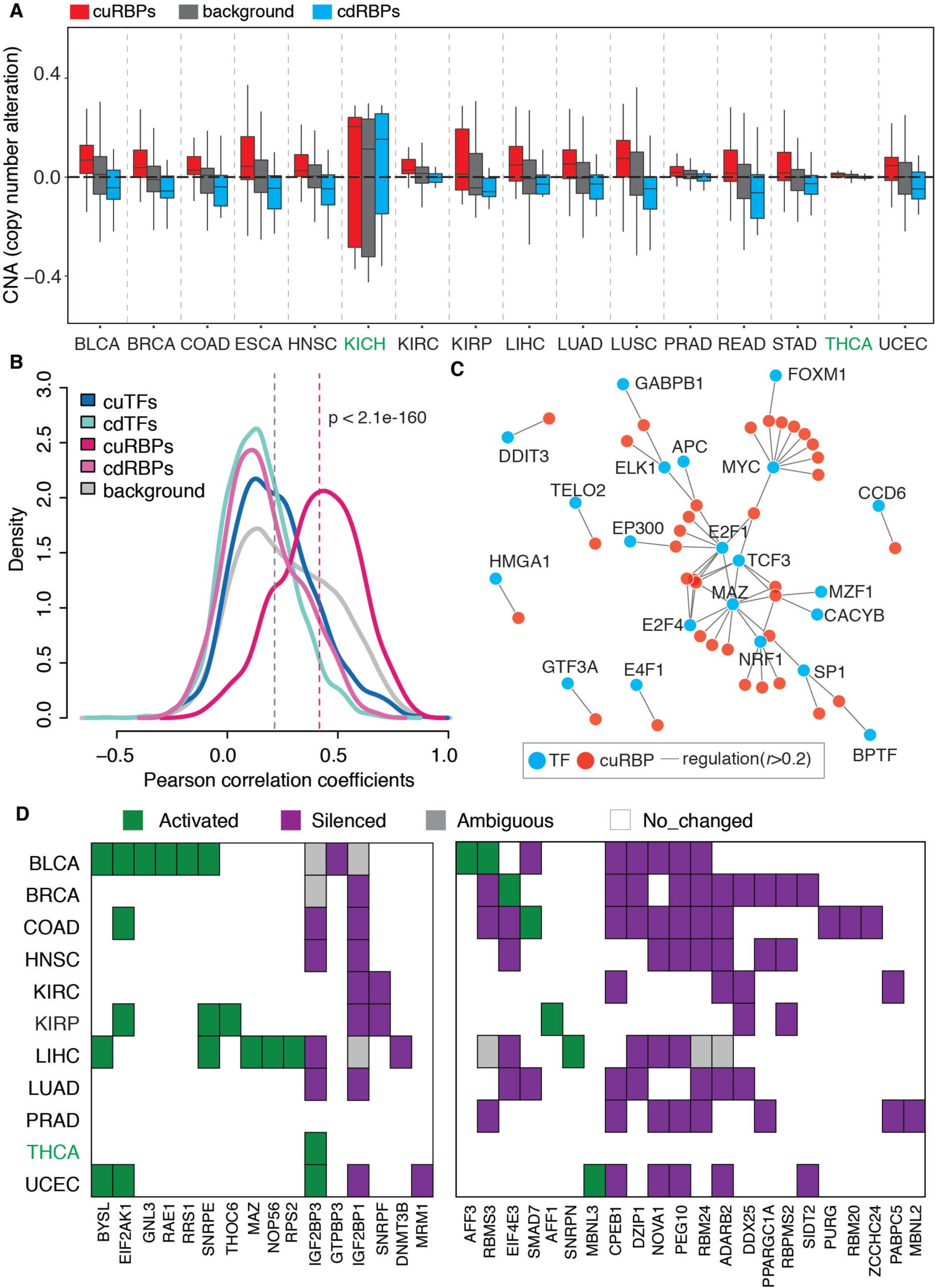
Potential mechanisms underlying dysregulation of the RBP expression in cancer. (A) The SCNA distribution of the cuRBPs, cdRBPs and background (all PCGs) across 16 cancer types in TCGA. The black horizontal dashed line indicates the median of the SCNA of all PCGs across 16 cohorts. (B) Density distribution of Pearson correlation coefficient between SCNA and gene expression for different gene groups. The *p* value represents the significance between cuRBPs and background. (C) Network between 21 positive regulatory TFs and their target cuRBPs (see Material and Methods). (D) Epigenetically activated, silenced and ambiguous cuRBPs and cdRBPs genes across 16 cancer types. Left panel are the 16 cuRBPs and right panel are 22 cdRBPs.

Besides changes in copy numbers, variation in the abundance of TFs such as MYC, and miRNAs could also result in dysregulated expression of RBPs (Ciafre and Galardi 2013; Koh et al. 2015). To identify the potential TFs and miRNAs which might be involved in the dysregulation of cuRBPs and cdRBPs in tumors, we utilized the gene sets (C3 sub-collection MIR: microRNA targets and C3 sub-collection TFT: transcription factor targets) from the GSEA molecular signature database (MSigDB), in which targets of TFs/miRNAs were predicted by searching the sequence motif either at the promoter region or within 3’ UTR respectively (Subramanian et al. 2005). In total, 500 TF and 221 miRNA gene sets were used for the analysis. For each TF-RBP or miRNA-RBP pair in each cancer type, we calculated the correlation between the expression of TF/miRNA and corresponding RBP. In total, 338 TF-cuRBP pairs, 265 TF-cdRBP pairs, 14 miRNA-cuRBP and 157 miRNA-cdRBP pairs were identified with correlation larger than 0.2 (TF: |*r*| > 0.2; miRNA: *r* < -0.2) in at least one cancer type. To find reliable TF/miRNA-RBP target relationships, we used more stringent criteria by requiring the correlation be consistent in more than 8 cancer types. Finally, 112 TF-cuRBPs pairs, 72 TF-cdRBPs and 16 miRNA-cdRBP pairs were obtained, while no miRNA-cuRBP pairs managed to fulfill this cutoff (Supplemental Table S3). For TFs that target cuRBPs, 21 of them are positive regulators (*r* > 0.2 in more than eight cancer types) and 29 are negative regulators (*r* < -0.2 in more than eight cancer types). We found that most of those TFs only target one to two cuRBPs, while MYC, MAZ, E2F1 and SP1 could target multiple RBPs (Figure 2C). Interestingly, SP1 is positively correlated with three cuRBPs and negatively correlated with 10 cdRBPs in our analysis. This finding is consistent with the previous view that SP1 is able to both enhance or repress promoter activity (Li et al. 2004).

DNA methylation has been shown to be an important epigenetic mechanisms for the regulation of gene expression (Jones 2012). To investigate whether the dysregulation of the RBPs is due to DNA methylation-mediated epigenetic alterations, we analyzed DNA methylation in 4607 tumors and 358 normal samples for around 0.3 million CpG sites among 11 cancer types based on the data in TCGA project. In general, we were unable to observe a significant decrease of DNA methylation at the promoters of cuRBPs, while DNA methylation at the promoters of cdRBPs were slightly elevated (Supplemental Fig. S4). Adapted from the method as described in previous study (Cancer Genome Atlas Research 2014), we classified the CpG sites into three groups: the epigenetically activated group, epigenetically silenced group and non-significantly changed group. The gene is defined as epigenetically activated or silenced if the promoter region contains either epigenetically activated or silenced CpG sites respectively. On the other hand, the gene is considered as ambiguous if the promoter region contains both epigenetically activated and silenced CpG sites. Using this classification, we identified 9 out of 109 (8%) cuRBPs to be epigenetically activated, while 22 out of 44 (50%) cdRBPs were epigenetically silenced in many cancer types. Moreover, despite being epigenetically silenced, 5 cuRBPs were observed to have significantly upregulated expression in tumors (Figure 2D).

The above results suggested that overexpression of cuRBPs in tumors is correlated with amplification of copy numbers, while downregulated expression of cdRBPs could be due to epigenetic silencing mediated by DNA methylation. Furthermore, some key regulators, such as MYC, MAZ, E2F1 and SP1 could also contribute to dysregulation of cuRBPs and cdRBPs in tumorigenesis.

### cuRBPs could work together to promote cancer progression while cdRBPs might function independently to repress tumorigenesis

To explore potential roles of the dysregulation of RBPs in cancer development, we investigated a series of their characteristics, including oncogenic/tumor suppressive potential, association with prognosis, connectivity between each other and the potential to form protein complexes. Based on the TUSON explorer, 300 tumor suppressor genes (TSGs) and 250 oncogenes (OGs) were identified in a previous study (Davoli et al. 2013). Among 1478 RBPs, 42 of them are potential TSGs and 22 of them are potential OGs, the proportion of TSG is higher than the background with statistical significance (Hypergeometric test, *p* < 0.0004). Next we asked whether the oncogenic RBPs are more likely to be upregulated, while tumor suppressive RBPs are more likely to be downregulated in cancer, by checking the numbers of TSGs and OGs in cuRBPs and cdRBPs. Unexpectedly, the results suggested that neither cuRBPs and cdRBPs have significantly more TSGs or OGs comparing with the background. However, there is slightly higher proportion of tumor suppressors in cdRBPs (Hypergeometric test, *p* < 0.1). As the TUSON explorer is based on the mutation pattern of genes across thousands of cancer patients, these data suggested that dysregulation of expression and mutation might contribute to cancer development independently, which is consistent with the view that alternative mechanisms of gene regulation could function distinctly in tumorigenesis (Santarius et al. 2010).

Recently established human pathology atlas provided the association between prognosis and expression of PCGs, in which 3755 favorable and 4476 unfavorable prognostic genes were identified (Uhlen et al. 2017). Interestingly, when comparing to TFs and PCGs, a higher proportion of RBPs are prognostically unfavorable (chi-squared test, *p* < 1.1e-18), in which increased expression is associated with poor overall survival rate. A more extreme trend is observed for cuRBPs, in which more than ~65% of cuRBPs are prognostically unfavorable while less than 5% are prognostically favorable. On the contrary, cdRBPs have a higher proportion of favorable prognostic genes than unfavorable ones (Figure 3A).

**Figure 3:**
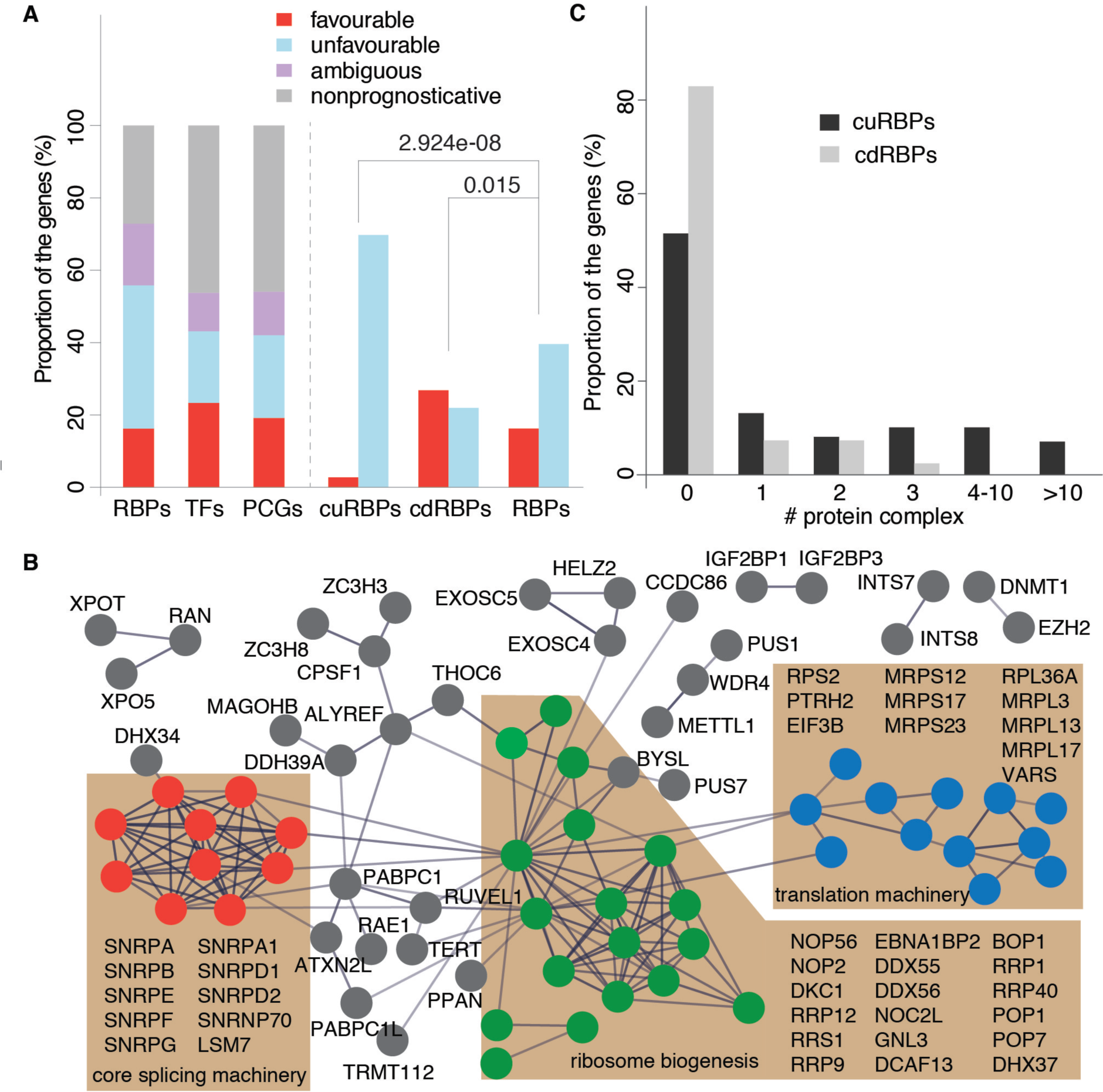
Characteristics of the dysregulated RBP in human cancers. (A) Distribution of the prognostic favorable, unfavorable and ambiguous RBPs. (B) PPI network of cuRBPs. Three most extensively connected modules are shown in red (splicing machinery), blue (translation machinery) and green (ribosome biogenesis) colors, while other proteins are shown in grey color. (C) Numbers of potential protein complexes formed by cuRBPs and cdRBPs.

Next, we checked the protein connectivity within cuRBPs, cdRBPs, cuTFs and cdTFs groups. Based on the protein-protein interaction (PPI) extracted from the STRING database, potential PPIs within each group were identified (Material and Methods) (Szklarczyk et al. 2017). Surprisingly the number of PPIs within cuRBPs was much higher than those in the other three groups and in fact, almost no PPI was observed within cdRBPs (Supplemental Fig. S5). This indicates that cuRBPs might function together, while cdRBPs might function independently. Three of the most extensively interacted modules in the PPI network of cuRBPs are: 1) core pre-mRNA splicing machinery in terms of small nuclear ribonucleoprotein (snRNP), 2) the translation machinery including ribosome components and translation initiation factors, and 3) the ribosome biogenesis regulators (Figure 3B). Besides the PPIs within each group, they might also form complexes with other proteins. To address this issue, we scanned through each RBP for its potential to form protein complexes based on the protein complex annotation from CORUM (Ruepp et al. 2010). We found that approximately half of the cuRBPs have the potential to form protein complexes, while more than 80% of the cdRBPs could not form any annotated protein complex (Figure 3C). The above results suggested that cuRBPs could work together to form protein complex including the core pre-mRNA splicing and translation machinery. Their overexpression might promote cancer progression as they were correlated with poor overall survival rate. On the other hand, cdRBPs might function independently to suppress tumorigenesis since there are almost no interactions among them and many of them are known tumor suppressors.

### Characterization of potential novel cancer-related RBPs in colon adenocarcinoma (COAD)

To investigate the functional consequences of the dysregulation of RBPs in tumorigenesis and identify potential oncogenic/tumor suppressive RBPs, we focused our analysis on colon adenocarcinoma (COAD). To confirm the general upregulation trend of RBPs in tumors, transcriptome analysis of a public COAD dataset (GSE104836), and an in-house microarray (Clariom) of 10 normal colon tissue versus 10 COAD tissue was carried out. Consistent with our findings in TCGA-COAD, both the RNA sequencing data from the public dataset and our microarray data show a much higher number of upregulated RBPs than downregulated ones (Figure 4A). As protein expression might not be correlated with mRNA expression, we next analyzed the protein expression in COAD and normal colon tissues from a previous study (Zhang et al. 2014). Again, we found that the protein expression of RBPs is significantly upregulated in tumors and this trend is even more notable for cuRBPs (Figure 4B).

**Figure 4:**
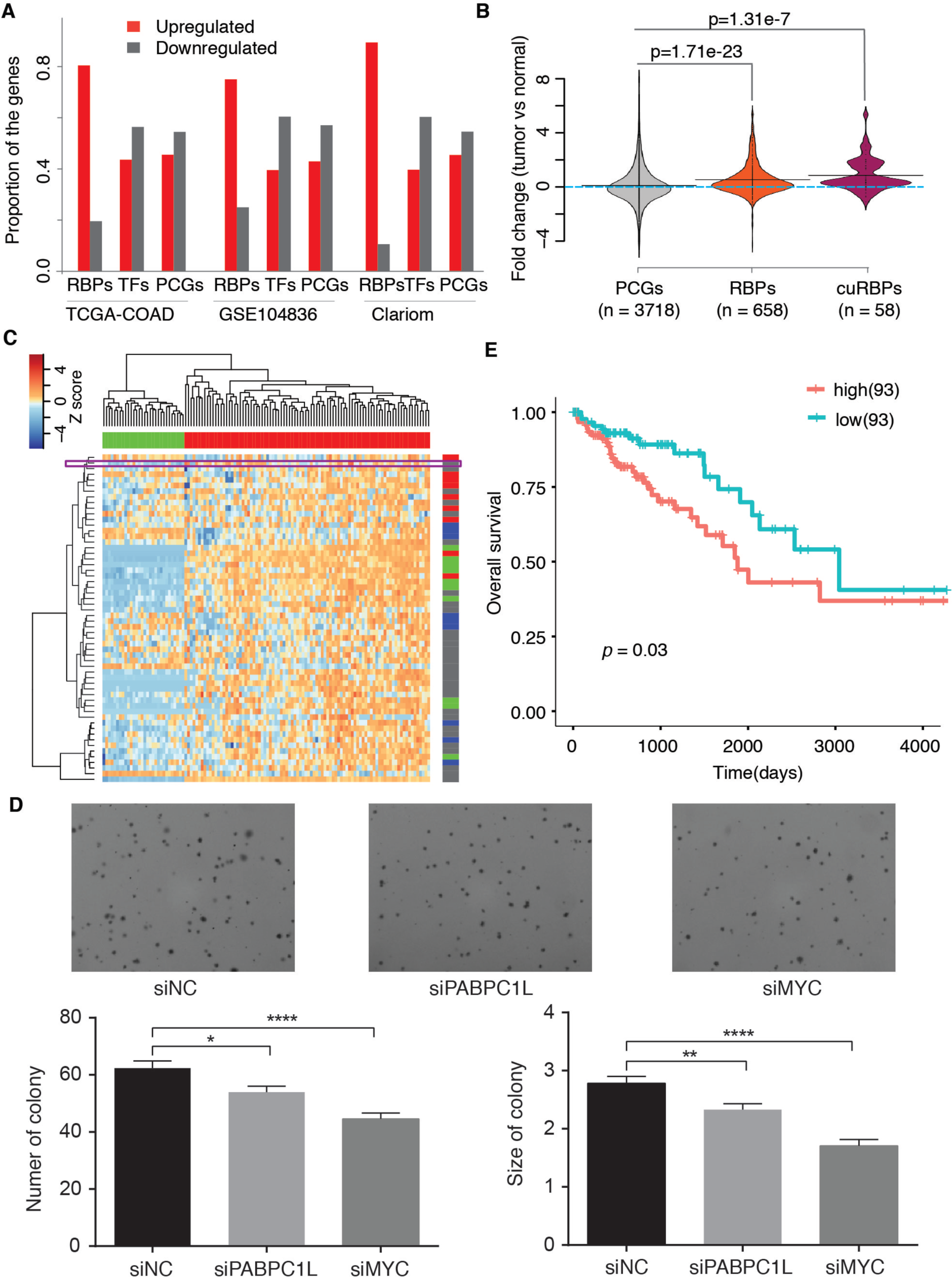
Characterization of potential novel cancer-related RBPs in COAD. (A) Proportion of dysregulated genes (tumor versus normal) in three independent datasets. TCGA-COAD: COAD dataset from TCGA project; GSE104836: public dataset containing the RNA sequencing data of 10 normal and 10 tumor COAD samples. Clariom: in-house microarray data for 10 COAD and 10 normal colon samples. (B) Distribution of the protein expression fold change between COAD tumor and normal samples. (C) Heatmap illustrating the protein expression of cuRBPs in normal and tumor samples. The horizontal bar: green stands for 30 normal samples and red for 91 tumor samples. The right vertical bar: red stands for the proteins related to splicing machinery, green for the proteins involved in ribosome biogenesis and the blue for the proteins related to translation machinery. The color and definition in the vertical bar is the same as Figure 3B. PABPC1L is highlighted in the purple box. (D) Soft agar assay of DLD cells upon PABPC1L, MYC knockdown. Error bar: SEM. t-test. *: p < 0.05, **: p < 0.01; ****: p < 0.0001. (E) Kaplan-Meier survival analysis of TCGA-COAD datasets based on segmentation values of PABPC1L expression.

It should be noted that hierarchical clustering of samples based on the protein expression profile of cuRBPs could distinctively separate tumor and normal, while proteins from the same functional module shared similar expression pattern and clustered together, such as the splicing machinery and proteins related to ribosome biogenesis (Figure 4C). To some extent these results further confirmed that the cuRBPs could function together to promote cancer progression by involving in core pre-mRNA splicing and translation machinery. Interestingly, we found several RBPs (RAE1, ALYREF, PPIH and PABPC1L) with similar protein expression to the RBPs of the core pre-mRNA splicing machinery, while three other RBPs, XPO5, XPOT and OAS3, were clustered together with the ribosome biogenesis related RBPs.

Among them, PABPC1L and RAE1 are frequently amplified in COAD patients (~10%) (Supplemental Fig. S6). Unlike RAE1 which is known to regulate transport of RNA between nucleus and cytosol, the function of PABPC1L is unclear. As its expression is correlated with spliceosome component, we speculate that it could mediate expression of core pre-mRNA splicing machinery RBPs such as SNRPD1, SNRPE and SNRPF. If so, the expression of PABPC1L should have a global effect on pre-mRNA splicing. To check this, we compared the splicing pattern between samples with high expression of PABPC1L (PABPC1L_high: 77 samples) and low expression of PABPC1L (PABPC1L_low: 77 samples). Indeed, we observed that 2502 significant events are more spliced in PBBPC1L_high group, while only 84 events are more spliced in PABPC1L_low group. This striking bias suggested that the PABPC1L could really function in RNA splicing (Supplemental Fig. S7).

Therefore, to further investigate the oncogenic function of PABPC1L, we performed soft-agar assay to assess its impact on cell proliferation in the COAD cell line DLD1. siRNA-mediated knockdown of the PABPC1L transcript resulted in a ~50% reduction in PABPC1L mRNA expression (Supplemental Fig. S8) and a concomitant, significant reduction in cell proliferation relative to the control (Figure 4D). Moreover, higher expression of PABPC1L is correlated with poor survival rate, suggesting that its dysregulation does contribute to cancer progression (Figure 4E).

## Discussion

In this study, starting with transcriptome analysis of RBP transcript expression in ~ 6700 samples across 16 cancer types, we observed an unexpected trend that RBPs are predominantly upregulated rather than downregulated in most cancer types. This is consistent with a previous study which observed the similar dysregulation pattern of RBPs in HCC, and is contrary to a recent study which reported a predominant downregulation instead of upregulation of RBPs across 15 human cancer types (Dang et al. 2017; Wang et al. 2018). By checking the expression of RBPs in more detail such as the heatmap and histogram of gene expression fold changes in some cancer types, as well as several public datasets and our in-house microarray data, we confirmed that RBPs are more likely to be upregulated in tumors. Interestingly, in both KICH and THCA, the number of upregulated RBPs is less than that of downregulated ones (Figure 1A). This suggested that both the KICH and THCA might behave differently from other cancer types and its underlying mechanisms need to be further explored.

In addition, we identified two groups of RBPs showing consistently up/down- regulation in almost all of the cancer types we studied. Many of them have been extensively studied for their roles in tumorigenesis, while the remaining few have unclear functional relevance to cancer. Thus our study not only provides insights to understand the dysregulation of RBPs during tumorigenesis, but also implicates several previously uncharacterized RBPs as potential oncogenes and tumor suppressors. Furthermore, we revealed distinct regulatory mechanisms underlying the dysregulation of cuRBPs and cdRBPs in tumors. The overexpression of cuRBPs is correlated with amplification of copy numbers, while the downregulation of cdRBPs could be due to DNA-methylation mediated epigenetic silencing (Figure 5).

**Figure 5:**
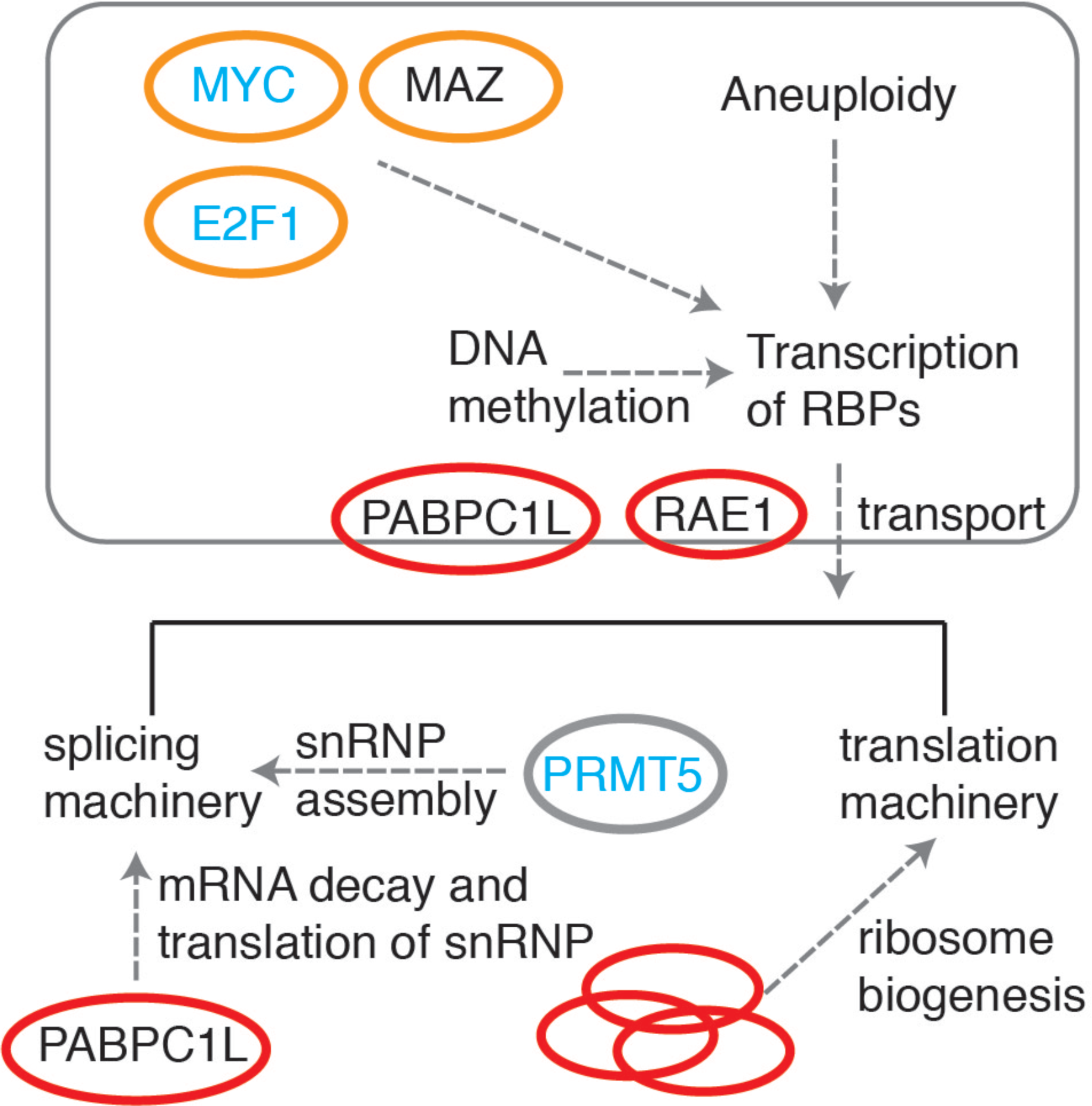
Potential role of the dysregulation of RBPs in cancer development.

Additionally, we also found that E2F1, MAZ, MYC and several other TFs could be upstream regulators that are responsible for dysregulation of RBPs in tumors (Figure 5). Surprisingly, according to the hallmark gene sets from the MSigDB (Liberzon et al. 2015), we identified 7 out of 50 hallmark gene sets where the RBPs are significantly enriched (Hypergeometric test by compared RBPs with background, BH-adjustment *p* < 0.05) and the most significant two are MYC_TARGET. Around 55% (33/58, BH-adjusted *p* < 1.1e-12) of the MYC_TARGET_V2 genes, 50% (101/200, BH-adjusted *p* < 8.7e-38) of the MYC_TARGET_V1 are RBPs (Supplemental Fig. S9). These results are consistent with previous studies which revealed that the expression of protein synthetic machinery such as ribosomal proteins, translation initiation factors, as well as core pre-mRNA splicing machinery, are essential for MYC mediated lymphomagenesis (Barna et al. 2008; Koh et al. 2015).

Despite the fact that more than half of the MYC_TARGET_V1 and MYC_TARGET_V2 genes are RBPs (125 RBPs in total), only 24 of them are cuRBPs. Among them, 9 cuRBPs might be regulated by MYC across many cancer types (Figure 2C and Supplementary Table 3). This is significantly higher than average (24/125 comparing to 109/1478, *p* < 0.0002, hypergeometric test), nevertheless it also suggested that the overexpression of more than 77% (85/109) of cuRBPs might be independent of MYC regulation. A recent study revealed that a specific RBP NELGF could selectively regulate downstream targets of MYC via stabilization of the transcript and its overexpression could enhance MYC mediated tumorigenesis (Dang et al. 2017). These results suggested that certain RBPs, particularly those which are genetically altered, may regulate critical functional processes and act as drivers of cancer progression.

A mode to elucidate the role of dysregulation of RBPs in tumors. Red ovals stand for RBPs and yellow ovals for TFs. The gene symbols in blue have known functions in tumorigenesis. The rectangular box indicates the nucleus.

Our results demonstrated that cuRBPs interact extensively with each other, with the core spliceosome, translational machinery and RBPs related to ribosome biogenesis as the most connected modules. This highlights that the overexpression of basic splicing and translation machinery could be crucial for cancer development. Indeed, inhibition of polymerase I (Pol I) that transcribes ribosome RNA (rRNA) by small molecular BMH-21, could repress tumor growth *in vivo* (Peltonen et al. 2014). On the other hand, depletion of the PRMT5 results in a decrease of the biogenesis of core pre-mRNA splicing machinery, repressing MYC induced lymphomagenesis in mice (Koh et al. 2015). A recent study revealed that ribosome levels has regulatory roles in hematopoietic cell differentiation (Khajuria et al. 2018). All these studies highlight that the abundance of basic splicing and translation machinery could be critical to maintain cell homeostasis and their dysregulation could promote tumorigenesis. Therefore, we hypothesized that the overexpression of RBPs which are components of basic splicing and translation machinery is required for MYC, E2F1, and other oncogene mediated tumorigenesis, while the regulators of the biogenesis of splicing and translation machinery might be critical to promote or even drive cancer progression (Figure 5).

Finally, our study found that the protein expression patterns of PABPC1L and RAE1 are similar to the core pre-mRNA splicing machinery components (Figure 4C). This could be attributed to several possible reasons: 1) they are co-regulated; 2), PABPC1L and RAE1 regulate the expression of splicing machinery components; 3), the core pre-mRNA splicing machinery regulates the expression of PABPC1L and RAE1. As the copy numbers of PABPC1L and RAE1 are frequently amplified in cancers (Supplemental Fig. 7), the most likely scenario is that PABPC1L and RAE1 regulate the expression of core pre-mRNA splicing machinery. Both these two proteins are from the RNA transport pathway, while PABPC1L is also involved in mRNA decay (Figure 5). Further studies should be performed to determine how they regulate the core pre-mRNA splicing machinery. Besides, we showed that the expression of PABPC1L is associated with mRNA splicing, cell proliferation and overall survival rate of COAD patients, suggesting that PABPC1L might promote cancer progression by regulating the core pre-mRNA splicing machinery. Therefore, dysregulation of PABPC1L might have similar functional consequences as PRMT5 and it could be a potentially valuable target for drug discovery to inhibit cancer progression as well.

## Materials and Methods

### Reagents

Reagents are as follows: siGENOME SMARTpool siRNA reagents (Dharmacon) for negative control (NC) (siNC), PABPC1L (siPABPC1L) and MYC (siMYC); DharmaFECT 1 transfection reagent (Dharmacon); Trizol reagent (ThermoFisher), Dulbecco’s modified Eagle medium (DMEM) (ThermoFisher), OptiMem reduced serum media (ThermoFisher), fetal bovine serum (FBS) (ThermoFisher), Trypsin-EDTA (ThermoFisher).

### Soft Agar assay

A 0.6% agarose base was prepared in 6-well dishes. At 24 h post-transfection, DLD-1 cells were trypsinized, re-suspended and counted. The cells (15 x 10^3^) were mixed with complete growth medium and agarose to a final agarose concentration of 0.3%, which was added above the base. The cells were grown at 37°C in a humidified atmosphere with 5% CO_2_ and were fed with complete growth medium every two days. After 8 days, the colonies were imaged under 4× magnification and quantified using ImageJ v.1.51k.

### Omics data and annotation resource

The RNA sequencing and the somatic copy number alteration (SCNA) as well as the DNA methylation data from TCGA project were downloaded and processed by TCGA-Assembler2 (Wei et al. 2017). In total, mRNA expression of 20530 genes in ~6700 clinical samples across 16 cancer types was obtained (Supplemental Table S1). The processed SCNA results contain the somatic copy number alteration of around ~16000 genes in each cancer in average. To exclude the impact of different platforms on the DNA methylation data, only the data from Infinium HumanMethylation450K BeadChip (Illumina) were used for the analysis, which include the normalized beta value for more than ~50 million CpG site across the genome.

Besides the data from the TCGA project, a public dataset containing 152 HCC and 91 normal liver samples (GSE102083) and another public dataset which contains 10 normal and 10 COAD tumor samples (GSE67526) were obtained from GEO. The protein expression data of 30 normal and 90 COAD tumor samples were obtained from a previous study (Zhang et al. 2014). The 1542 RBPs were defined from RBP census (Gerstberger et al. 2014), while only 1478 of them have available expression data in the TCGA datasets. 1290 TFs were derived from DBD human transcription factor database (Wilson et al. 2008). The oncogenes and tumor suppressor genes were defined based on the TUSON explorer score as described in a previous study (Davoli et al. 2013). In brief, 18679 genes were ranked by q value, the top 300 with smallest TUSON_q_value_TSG and top 250 with smallest TUSON_q_value_OG were selected as oncogenes and tumor suppressors respectively. 6114 prognostic favorable genes and 6834 unfavorable genes were obtained from the human pathology atlas (Uhlen et al. 2017). Therein, 2357 genes that could be observed in both prognostic favorable and unfavorable lists were defined as ambiguous. 3756 prognostic favorable genes and 4476 unfavorable genes were defined by removing those ambiguous ones. The genetic alteration of seven candidate RBPs were obtained from cBioPortal (Gao et al. 2013).

### Transcriptome data analysis

As TCGA datasets we studied contain at least 10 normal and many more tumor samples (usually hundreds of samples) in each cancer cohort (Supplemental Table S1), we performed the t-test to evaluate expression difference between tumor and normal samples in each cancer cohort and used Benjamin-Hochberg (BH) approach to adjust the p value similar to previous study (Dang et al. 2017). The genes with BH-adjusted p value smaller than 0.001 were considered as dysregulated genes. For each cancer type, the ratio between upregulated RBPs and downregulated RBPs was compared to that of TF and background that contains all the 20531 PCGs. The significance of the difference between RBPs and TFs, RBPs and background were estimated by hypergeometric test. The consistently up/down-regulated RBPs and TFs were defined based on two metrics: the directionality and amplitude. For each gene, the directionality was the difference between the number of cancer types, in which the gene is upregulated and the number of cancer cohorts, in which the gene is downregulated. The amplitude was the average of the expression fold change across 16 cancer types. The RBPs with directionality larger than 8 and amplitude larger than log_2_(1.5) was defined as cuRBPs, while those with directionality smaller than -8 and amplitude smaller than –log_2_(1.5) were defined as cdRBPs. In total, 109 cuRBPs and 41 cdRBPs were obtained. Similarly, 45 cuTFs and 137 cdTFs were defined by the same criteria. All of the heatmaps of gene expression in normal and tumor samples were generated by R function heatmap.2 from “gplots” package.

### Microarray preparation

RNA quality was assessed by using the Agilent Model 2100 Bioanalyzer (Agilent Technologies, Palo Alto, CA). 150nanogram of total RNA was processed for use on the microarray by using the Affymetrix WT plus kit according to the manufacturer’s recommended protocols. The resulting biotinylated cRNA was fragmented and then hybridized to the Clariom D array (Applied Biosystems). The arrays were washed, stained, and scanned using the Affymetrix Model 450 Fluidics Station and Affymetrix Model 3000 7G scanner using the manufacturer’s recommended protocols by the Microarray Facility. Expression values were generated by using Expression Console software (Affymetrix). Each sample and hybridization underwent a quality control evaluation.

### SCNA analysis

Based on the SCNA data from TCGA, copy number alterations of cuRBPs and cdRBPs were compared to the corresponding background of all the PCGs with Mann-Whitney-Wilcoxon test for each cancer type. In addition, the Pearson correlation coefficient between the gene expression (log transformed normalized RSEM value) and SCNA were calculated for each gene in a group. The significance of difference of the correlation coefficients between the gene sets (cuRBPs/cdRBPs/cuTFs/cdTFs) and background (PCGs) were estimate by MannWhitney-Wilcoxon test.

### TF/miRNA analysis

To identify potential TF/miRNA regulators of RBPs, 500 gene sets of TFs, 221 gene set of miRNAs were obtained from molecular signature database (MSigDB, version 6), in which targets of TFs/miRNAs were predicted based on the motif at the promoters/3’ UTRs. To determine whether dysregulation of a TF/miRNA in tumor is really functional to its target cuRBPs or cdRBPs, we calculated the correlation in expression between this TF/miRNA and each of its target cuRBP and cdRBP in each cancer type separately. The TF-cuRBP and TF-cdRBP pairs with correlation larger than 0.2 in more than eight cancer types were defined as positive regulation, while correlation smaller than -0.2 were defined as negative regulation. For the miRNA-cuRBP and miRNA-cdRBP pairs, the correlation smaller than -0.2 in more than eight cancer types were required.

### DNA methylation analysis

To investigate the link between DNA methylation and transcription of a gene, only the CpG sites at its promoter region were selected. Based on the GENCODE (Version 27 liftover to hg19) annotation and definition from the DNA methylation database (Huang et al. 2015), 5’ UTR and 1700bp (TSS200 and TSS1500) upstream region of transcription start site (Uhlen et al.) of a gene were defined as promoter. By comparing normalized DNA methylation values (β) between tumor and normal samples, the epigenetically silenced and activated CpG sites were determined with a similar approach as described in the TCGA project. The epigenetically silenced CpG site must fit the following three criteria: 1), more than 90% of normal samples are un-methylated (β < 0.1); 2) the difference of DNA methylation average between tumor and normal is larger than 0.2; 3) The difference between tumor and normal should be significant (BH-adjusted *p* < 0.05). The epigenetically activated CpG sites were identified by a similar approach but the change is in the opposite way: 1), at least 90% of normal samples is methylated (β > 0.3), 2) the difference is smaller than -0.2; 3) BH-adjusted *p* < 0.05. Those genes, whose promoter only contains epigenetically activated/silenced CpG site were defined as epigenetically activated/silenced. If the promoter of a gene contains both the epigenetically silenced and activated CpG sites, it was defined as ambiguous.

### Protein-protein interactions and protein complexes

The putative protein-protein interactions (PPIs) within each groups, including the cuRBPs, cdRBPs, cuTFs and cdTFs were identified based on the STRING database (version 10.5) (Szklarczyk et al. 2017). The medium confidence with interaction score > 0.4 was set as cutoff for an existing interaction between two proteins and only the interactions with experimental evidence were used. To evaluate the capacity of each protein to form protein complexes, the complex annotation was downloaded from the CORUM database (Ruepp et al. 2010). To compare the connectivity within each group, the ratio of the number of nodes (proteins) and edges (PPIs) was assessed and the significance between different groups was estimated by hypergeometric test.

## Data access

Two public GEO datasets including the GSE102083 and the GSE67526 (https://www.ncbi.nlm.nih.gov/geo/) were used in this study. The Clariom microarray was deposited to GEO as well under accession number (GSE115261).

## Acknowledgements

We thank all members in Tay lab and Henry lab for critical comments and suggestions. We thank Parul Saxena for microarray service and National Supercomputing Centre Singapore (NSCC) for computing service. Y.T. was supported by a Singapore National Research Foundation Fellowship, a National University of Singapore President’s Assistant Professorship and the RNA Biology Center at CSI Singapore, NUS, from funding by the Singapore Ministry of Education’s Tier 3 grant number MOE2014-T3-1-006.

## Author contributions

Y.T and H.Y conceived and led the project and B.Z performed the data analysis and wrote the manuscript. K.R.B and C.Y.L did the soft-agar assay experiment and K.Z.H performed microarray experiment. J.L provided splicing results and S.Z and K.R.B helped to modify the manuscript.

